# An NHEJ-independent Role for DNA-PKcs in ATR activation at DNA Double-Strand Breaks

**DOI:** 10.64898/2026.06.19.733426

**Authors:** Oanh Huynh, W. Matthew Michael

## Abstract

DNA double-strand breaks (DSBs) are threats to genome integrity, and to mitigate this risk cells activate the Ataxia Telangiectasia and Rad3-related (ATR) kinase, which halts cell cycle progression to allow time for repair. While ATR signalling during replication stress is well understood, how ATR is activated at DSBs remain unclear. Topoisomerase 2-Binding Protein 1 (TOPBP1) is a key activator of ATR, and activation is mediated by phosphorylation of TOPBP1 at Serine 1131 (S1131). Previous work showed that the Ataxia Telangiectasia Mutated (ATM) kinase phosphorylates TOPBP1 at S1131. ATM is primarily linked to the homologous recombination (HR)-based repair of DSBs, however the majority of cellular DSBs are repaired via the Non-Homologous End-Joining (NHEJ) repair pathway, raising the question of how (or if) ATR is activated in an ATM-independent manner. Here, using *Xenopus* egg extracts, we demonstrate that DNA-PKcs controls a pathway acting in parallel to ATM that promotes ATR signalling at DSBs. We show that, like ATM, DNA-PKcs phosphorylates TOPBP1 at S1131. DNA-PKcs is best known for orchestrating NHEJ, however we find that its roles in NHEJ and ATR signalling are separable. Our findings reveal an alternative pathway for ATR activation in which DNA-PKcs directly couples DSB recognition to ATR signalling.

**GRAPHICAL ABSTRACT:** 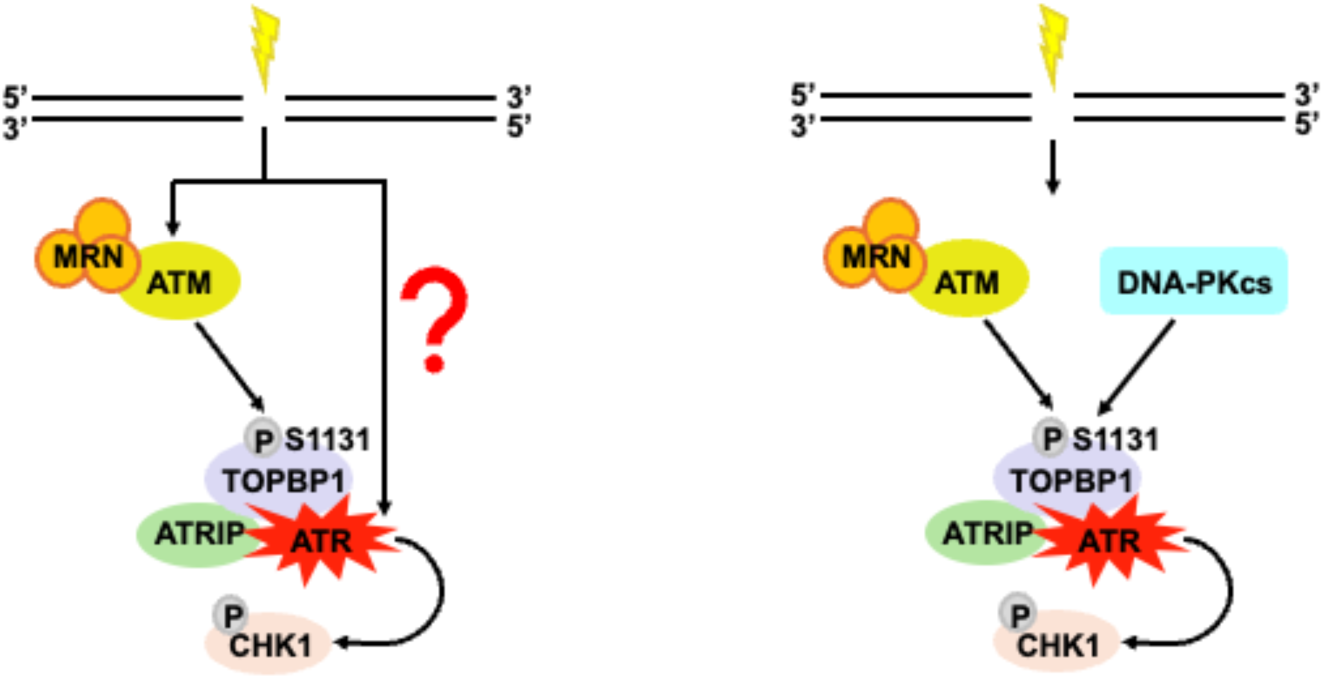

## INTRODUCTION

DNA double-strand breaks (DSBs) are the most dangerous form of DNA damage because they can lead to genome instability and chromosome rearrangement if not detected or repaired properly. To maintain genome integrity and cellular homeostasis, cells activate DNA damage response pathways orchestrated by a family of phosphatidylinositol 3-kinase-related kinases (PIKKs) that include DNA-Dependent Protein Kinase catalytic subunit (DNA-PKcs), Ataxia Telangiectasia Mutated (ATM), and Ataxia Telangiectasia and Rad3-related (ATR) kinases [1]. Once activated, ATM and ATR phosphorylate hundreds of substrates, including Checkpoint Kinase 2 (CHK2) and Checkpoint Kinase 1 (CHK1), respectively, to halt cell cycle progression, induce apoptosis, and facilitate DNA repair [2]. While extensive studies have characterized the roles of DNA-PKcs and ATM in responding to DSBs, the mechanism by which ATR is activated at DSBs remains unclear.

In parallel with DSB sensing, cells activate one of two major DNA repair pathways - non-homologous end-joining (NHEJ) or homologous recombination (HR). NHEJ is an error-prone pathway and is active throughout the cell cycle, but predominantly in the G1 and G2 phases [3, 4]. In the NHEJ pathway, the Ku heterodimer, consisting of Ku70 and Ku80, rapidly binds to the ends of DSBs [5]. Subsequently, the Ku complex recruits DNA-PKcs along with XLF and the XRCC4–LIG4 complex to facilitate repair [6]. By contrast, HR is an error-free pathway as it uses a sister chromatid as a template for strand invasion and recombination and functions in the S-phase and also plays a minor role in G2 [3,4]. In the HR pathway, the MRN complex, consisting of <u>M</u>RE11, <u>R</u>AD50, and <u>N</u>BS1 subunits, binds and recruits ATM to DSBs. MRN, coupled with ATM, initiates DNA end resection that results in RPA-coated ssDNA, allowing strand-invasion and recombination to proceed [7].

The prevailing model of ATR activation at DSBs is that DNA end resection creates a tract of Replication Protein A (RPA) coated ssDNA that allows recruitment of ATR and its binding partner ATR-Interacting Protein (ATRIP) [8]. In addition, ATM is recruited to DSBs and activated via interaction with its partner, the MRN complex [9, 10]. ATR recruitment is not sufficient for its activation, however, as activation requires another factor, the TOPBP1 protein [11, 12, 13]. TOPBP1 binds both ATRIP and ATR in a manner that stimulates ATR catalytic activity [11, 14]. Previous work has shown that ATM phosphorylates TOPBP1 on S1131, to create P-S1131, and that this modification is required for TOPBP1 to then activate ATR [12, 15] Thus, for HR-dependent repair of DSBs, a model has emerged where ATR activation is dependent on ATM via the requirement for P-S1131 on TOPBP1.

Over the past several years, we have employed an experimental system based on *Xenopus* egg extracts to study how ATR is activated by DNA DSBs. In this system, termed DMAX for <u>D</u>SB-<u>m</u>ediated <u>A</u>TR activation in *<u>X</u>enopus* egg extracts, we use linearized plasmid DNAs as a source of DSBs and have shown that ATR signaling in DMAX is absolutely dependent on TOPBP1, but only partially dependent on MRN and ATM [13, 16]. Indeed, when MRN is removed from the extract, we observed a decrease, but not an elimination, of ATR signaling, and the same was true when ATM was inhibited using a highly specific chemical inhibitor [16]. These data suggest that, at least in *Xenopus*, multiple DSB-dependent ATR activation pathways are operational. MRN and ATM-independent mechanisms for ATR signaling are also consistent with the fact that most cellular DSBs are repaired via NHEJ [3], which does not absolutely rely on end resection for repair. As such, the challenge now is to reconcile the absolute requirement for TOPBP1 P-S1131 with the partial requirement for MRN/ATM in DSB-induced ATR signaling.

In this study, we identify a Ku-independent role for DNA-PKcs in ATR activation at DSBs that operates in parallel with ATM. We demonstrate that while Ku70/80 and XLF are essential for NHEJ, they are dispensable for ATR signaling. By contrast, DNA-PKcs kinase activity is required for ATR activation, and DNA-PKcs can associate with DSBs independently of the Ku complex. Mechanistically, we provide evidence that DNA-PKcs phosphorylates TOPBP1 at S1131 and functions in parallel with ATM to promote ATR activation. These findings reveal an alternative signaling pathway in which DNA-PKcs links DSB recognition to ATR activation, thereby expanding the current model of DSB-induced ATR signaling.

## MATERIAL AND METHODS

### Plasmids

| Plasmid | Name | Vector | AA Coordinates |
| --- | --- | --- | --- |
| Myc-TOPBP1 | Cut5 | pCS2+MT | 1 – 1513 |
| Myc-TOPBP1-S1131A | pSY29 | pCS2+MT | 1 – 1513 (S1131A) |
| Myc-TOPBP1-S1131D | pKM91 | pCS2+MT | 1 – 1513 (S1131D) |
| XLF | pOH78 | pCS2 | 1 – 297 |
| Ku70 | HGNJSZ_1_3.1* | pCS2 | 1 - 721 |
| Ku80 | HGNJSZ_4_44.1* | pCMV-SPORT6 | 1 - 818 |
| GST (*E.coli* expression vector) | - | pGEX-4T3 | - |
| GST-AAD (*E. coli* expression vector) | pHG127 | pGEX-4T3 | 972 -1279 |
| GST-AAD-S1131A (*E.coli* expression vector) | pKR10 | pGEX-4T3 | 972 -1279 (S1131A) |
| GST-AAD-S1131D (*E.coli* expression vector) | pKR11 | pGEX-4T3 | 972 -1279 (S1131D) |
| DNA-PKcs (*E.coli* expression vector) | pTG314** | pET28a | 3765 - 4146 |

\*: Kind gift of Hironori Funabiki, Rockefeller University)
\*\* : Kind gift of Joseph Loparo, Harvard University

The *Xenopus* XLF DNA sequence, which was retrieved from [17], was synthesized by TwistBioscience and subcloned into the pCS2 vector (Forward primer: 5’ATGGATGCCCGACTCTTGCAGCTGCCC3’; reverse primer: 5’GAATTCTTACATAAACAGTCCCTTTGCCTTCTTTTT 3’)

### Chemical inhibitors

| Inhibitor | Name | Source | Catalog Number |
| --- | --- | --- | --- |
| DNA-PKcs | NU7441 | Selleckchem | S2638 |
| DNA-PKcs | AZD7648 | MedChemExpress | HY-111783 |
| ATM | KU60019 | Selleckchem | S1570 |
| ATR | Compound 45 | Charrier et al., 2011 |  |
| MRE11 | Mirin | Sigma | M9948-5MG |

### Recombinant proteins

The GST and GST-AAD recombinant proteins were purified as follows. The plasmids were expressed in *Escherichia coli* BL21 (DE3) at 37°C for 3 hours, then induced with 1 mM IPTG at 16°C overnight. The soluble proteins were then purified according to standard procedures. For antibody production, *Xenopus* 6His-DNA-PKcs (plasmid pTG314) was expressed in *E. coli* BL21 (DE3) at 37°C for 3 hours, then induced with 1 mM IPTG at 16°C overnight. The insoluble protein was purified under denaturing conditions using a procedure adapted from the Qiagen manual for purification of 6×His-tagged proteins from *E. coli*.

### Antibody production

The *Xenopus* peptides of Ku80 (CMEDEGDVDDLLDMM; sequence retrieved from [19]) and XLF (CGASKPKKKAKGLFM; sequence retrieved from ref. 17), with KLH conjugation on cysteine, were synthesized by GenScript. Anti-Ku80 and anti-XLF antibodies were raised in rabbits against the synthetic peptides, and anti-DNA-PKcs antibody was raised in rabbits against *Xenopus* 6His-DNA-PKcs by Pocono Rabbit Farm and Laboratory. The Ku80, DNA-PKcs, and XLF antibodies were affinity-purified using the AminoLink™ Plus Immobilization Kit (Thermo Scientific, 44894).

### Antibodies

The following antibodies were purchased from commercial suppliers: GST (Millipore Sigma #05-782), CHK1 (Santa Cruz Biotechnology #sc-8408), phosphorylated CHK1 (P-CHK1) (Cell Signaling Technology #2341S), and phosphorylated ATM S1981 (P-ATM) (Rockland #200-301-400S). Antibodies against *Xenopus* Ku80, DNA-PKcs, and XLF were produced by us in collaboration with Pocono Rabbit Farm and Laboratory. The DNA-PKcs 4227 antibody, used to detect DNA-PKcs by Western blotting, was a kind gift from Michael Lieber (University of Southern California). The *Xenopus* anti-p-S1131 TOPBP1 (P-TOPBP1) antibody was a kind gift from William Dunphy (California Institute of Technology). DNA-PKcs antibody was used for immunoprecipitation, and DNA-PKcs 4227 was used for detection by Western blotting.

### High-speed supernatant (HSS) *Xenopus* egg extracts

The high-speed supernatant (HSS) of *Xenopus* egg extracts, hereafter referred to as XEE, was prepared as described in [21] and used exclusively in this work.

### **I**n <u>v</u>itro transcription/translation (IVTT) of recombinant proteins

IVTT reactions were performed using the SP6 TnT Quick Coupled Transcription/Translation System (Promega #L2080) according to the manufacturer’s instructions. The recombinant proteins were used directly without downstream purification.

### Immunodepletion

For TOPBP1 immunodepletion, the HU142 antibody was used, and the procedure was performed as described by [21]. For immunodepletion of the Ku complex (via anti-Ku80) and XLF, the Ku80 and XLF antibodies were used, respectively, following the same procedure as for TOPBP1 immunodepletion. Depleted XEE was supplemented with IVTT recombinant proteins for rescue conditions.

### Immunoprecipitation

The immunoprecipitation procedure was adapted from [18]. Briefly, 20 µl of HSS XEE was incubated with protein A beads (Cytiva, 17127901) coupled to TOPBP1 (HU142) or DNA-PKcs antibody for 1 hour at 4°C. The beads were then washed three times with Buffer A (10 mM HEPES-KOH, pH 7.5, 150 mM NaCl, 0.5% Nonidet P-40, 2.5 mM EGTA, and 20 mM β-glycerophosphate) and twice with HEPES-buffered saline (10 mM HEPES-KOH, pH 7.5, 150 mM NaCl). The beads were eluted in 2X Laemmli sample buffer, and the samples were subjected to Western blotting.

### Kinase assay

The kinase assay procedure was adapted from [12, 18]. 50 µl of mock-depleted XEE or XEE depleted of endogenous TOPBP1 was incubated with Protein A beads (Cytiva, 17127901) coupled to a DNA-PKcs antibody for 1 hour at 4°C. The beads were washed three times with 80 mM β-glycerophosphate (pH 7.3), 20 mM EGTA, 15 mM MgCl2, and 1 mM dithiothreitol and twice with HEPES-buffered saline (10 mM HEPES-KOH, pH 7.5, and 150 mM NaCl). The beads were resuspended in kinase buffer (50 mM Tris-HCl [pH 7.5], 10 mM MgCl2, 1 mM dithiothreitol, and 1 mM ATP) and incubated with unprogrammed or programmed IVTT TOPBP1 for 30 minutes at room temperature. The reactions were stopped by adding 2X Laemmli sample buffer and boiling. The samples were subjected to Western blotting.

### End-joining assay

20 µl of HSS XEE was combined with 0.4 µl energy mixture (EM; 375 mM creatine phosphate, 50 mM ATP pH 7.4, 5 mM EGTA pH 7.7, 50 mM MgCl₂) and DSB substrate (AfeI-digested 8.9 kb plasmid) at a final concentration of 15 ng/µl). DMSO or chemical inhibitors were added to the reaction mixture as applicable. Reactions were incubated for the indicated time points and stopped by adding stop buffer (2% SDS, 0.5 M EDTA pH 8.0), followed by digestion with RNase A (10 mg/ml, Fisher Scientific) and Proteinase K (800 U/ml, NEB). Samples were separated on a 1× TAE 0.6% agarose gel and post-stained with SYBR Green (Thermo Fisher, S7567).

### DMAX assay

20 µl of HSS XEE was mixed with okadaic acid (1 µM). DMSO or chemical inhibitors were added to the reaction mixture as applicable. DSB substrate (AfeI-digested 8.9 kb plasmid) was added to the XEE mixture, and the reactions were incubated at room temperature for the indicated time points. Samples were collected, mixed with 2X Laemmli sample buffer, and analyzed by Western blotting.

### DSB-binding assay

A biotinylated 8.9-kb PCR fragment was generated using a forward primer bearing a biotin moiety. The fragment was coupled to magnetic streptavidin beads (Dynabeads™ Streptavidin Magnetic Beads, Thermo Fisher Scientific) according to the manufacturer’s instructions. These “DSB beads” contained 600 fmol of dsDNA per assay. 20 µl of mock-depleted XEE or XEE depleted of the protein of interest was supplemented with unprogrammed or programmed IVTT proteins. The XEE mixture was incubated with the DSB beads for 30 minutes. In Figure S2B, no XEE was used — only IVTT lysates. Samples were placed on a magnetic stand. The bound fraction was collected and washed three times in PBS + 0.1% Triton X-100, eluted with 2X Laemmli sample buffer, and analyzed by Western blotting.

### Data analysis

Signal intensities of proteins were quantified using ImageJ. The positive control was set to 100, and the remaining conditions were adjusted accordingly. Experiments shown in Figs. 1A–1C and 6D were performed in triplicate; graphs were plotted from the triplicate data, error bars represent standard deviations, and p-values were calculated using a t-test (n.s., not significant, p > 0.05; *, p < 0.05; **, p < 0.0005; ***, p < 0.00005). Two biological replicates were performed for each of the remaining experiments.

**Figure 1:**
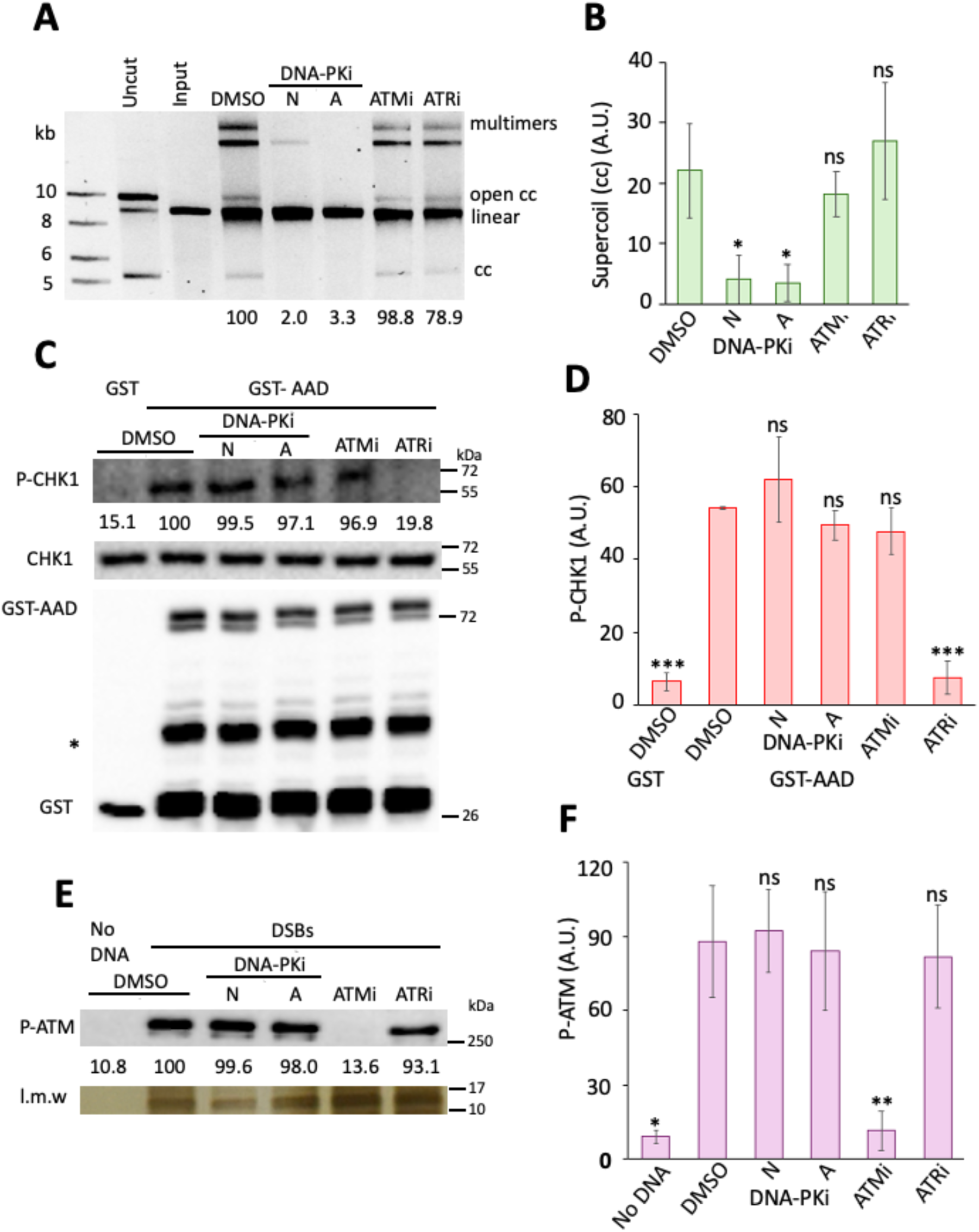
Validation of small-molecule inhibitors used in this study. A, a representative end-joining assay assessing NHEJ repair efficiency to validate inhibitors targeting DNA-PKcs, ATR, or ATM. XEE was treated with a vehicle-only control (DMSO), a DNA-PKcs inhibitor (DNA-PKi: 100 µM NU7441 [N] or 10 µM AZD7648 [A]), an ATM inhibitor (ATMi: 100 µM KU60019), or an ATR inhibitor (ATRi: 100 µM Compound 45). The XEE mixture was combined with the energy mix and the DSB substrate. After 10, 20, or 30 minutes of incubation, samples were digested with RNase A and Proteinase K and subjected to 0.6% agarose gel electrophoresis, followed by post-staining with SYBR Green. Three biological replicates were performed, and a representative experiment is shown. Supercoil (cc) signal intensity was quantified using ImageJ; the signal from the DMSO-treated sample was set to 100, and all other samples were normalized accordingly. B, a bar graph showing quantification of the raw arbitrary units (A.U.) of supercoils. C, a representative assay assessing ATR activation to validate inhibitors targeting DNA-PKcs, ATR, or ATM. XEE was treated with DMSO, DNA-PKi (100 µM NU7441 [N] or 10 µM AZD7648 [A]), ATMi (100 µM KU60019), or ATRi (100 µM Compound 45). Subsequently, GST or GST-AAD was added to the XEE mixture. All samples were incubated for 10, 20, or 30 minutes. After incubation, samples were subjected to Western blot analysis. Three biological replicates were performed, and a representative experiment is shown. Total CHK1 served as a loading control. P-CHK1 signal intensity was quantified using ImageJ; the P-CHK1 signal from the DMSO-treated sample was set to 100, and all other samples were normalized accordingly. D, a bar graph showing quantification of the raw arbitrary units (A.U.) of P-CHK1. E, a representative DSB-binding assay determining ATM autophosphorylation to validate inhibitors targeting DNA-PKcs, ATR, or ATM. XEE was treated with DMSO, DNA-PKi (100 µM NU7441 [N] or 10 µM AZD7648 [A]), ATMi (100 µM KU60019), or ATRi (100 µM Compound 45). “Input” refers to a sample of the total XEE taken before the addition of the DSB beads. “Bound” refers to proteins bound to the DSB beads. Three biological replicates were performed, and a representative experiment is shown. A low-molecular-weight (l.m.w.) band, likely histone, served as a loading control and was visualized by silver staining. P-ATM signal intensity was quantified using ImageJ; the P-ATM signal from the DMSO-treated sample was set to 100, and all other samples were normalized accordingly. F, a bar graph showing quantification of the raw arbitrary units (A.U.) of P-ATM. In panels B, D, and F, error bars represent standard deviations; p-values were calculated using a t-test (n.s., not significant, p > 0.05; *, p < 0.05; **, p < 0.0005; ***, p < 0.00005). Uncut, circular 8.9 kb plasmid; Input, AfeI-digested 8.9 kb plasmid, linear; DNA repair products: multimers, open circles, and closed circles (cc; supercoils).

## RESULTS

### Validation of the PIKK inhibitors used in this study

The goal of our study was to determine the relationship between NHEJ-based DNA repair and ATR signaling at DSBs, which necessitated an analysis of the role of PIKK family members ATM and DNA-PKcs in activation of ATR kinase by DSBs. In recent years, highly specific chemical inhibitors have been developed that target ATM, ATR, and DNA-PKcs, and as these are attractive reagents for our project, it was important to first validate their specificity in our DMAX system. To evaluate DNA-PKcs activity, we used a simple end-joining assay, where a linearized and blunt-ended plasmid DNA of 8.9 kb was incubated in the high-speed supernatant (HSS) produced after centrifugation of crude egg extract at 100,000 x *g*. The high-speed spin removes membrane vesicles and endogenous DNA from the extract and thus HSS is composed of soluble proteins, which are present at a concentration of 50 mg/ml. As seen in Fig. 1A, incubation of the linearized dsDNA in HSS treated with vehicle (DMSO) led to a conversion of the linear DNA to either a closed circle (via intramolecular end joining) or to multimers (dimers and trimers, via intermolecular end joining) after 30 minutes of incubation. Previous work, using an identical extract system, had shown that addition of 100 µM NU7441, an inhibitor of DNA-PKcs, blocks end joining, and we observed the same effect in our experiment (Fig. 1A&B). We note that work with egg extracts often requires chemical inhibitor concentrations that are 10–100 times higher than those used in standard cell culture or purified protein assays [20]. This necessity is driven primarily by the exceptionally high protein concentration of the extract, which creates several biochemical challenges such as high target concentration or, more commonly, nonspecific sequestration of the compound by the crowded molecular environment. Recently, a next-generation DNA-PKcs inhibitor has been developed, AZD7648 [21], and we found that it too would efficiently block end-joining in HSS when employed at concentrations as low as 10 µM (Fig. S1 and Fig. 1A&B). For the experiments presented below, we therefore used both NU7441 and AZD7648 to inhibit DNA-PKcs.

We next asked if addition of 100 µM KU60019, which inhibits ATM (referred to as ATMi, [22]) or 100 µM Compound 45, which inhibits ATR (referred to as ATRi, [23]), would have an impact on end joining and found that they did not (Fig. 1A&B). These data show that neither ATMi nor ATRi can “spill over” and inhibit DNA-PKcs in our system. To pursue these observations, we next asked if the DNA-PKcs inhibitors, or ATMi, could spill over and target ATR in our system. To assess ATR signaling, we utilized a recombinant protein, GST fused to the TOPBP1 ATR activation domain (AAD), to activate ATR. Previous work has shown that addition of GST-AAD to HSS robustly activates ATR [11, 24]. To monitor ATR activity, we used Western blotting and an antibody that recognizes the serine 345 phosphorylated form of CHK1 (P-CHK1), a critical ATR substrate. We also probed for unmodified CHK1, as a loading control. As shown in Fig. 1B and quantified in Fig. 1C, addition of GST-AAD, but not GST alone, led to the appearance of P-CHK1 in control samples. Importantly, P-CHK1 was greatly reduced in samples containing ATRi, as expected, but was not impacted by either DNA-PKcs inhibitor or ATMi (Fig. 1C&D). Thus, neither DNA-PKcs inhibitor nor ATMi can block ATR kinase activity. Lastly, we asked if the DNA-PKcs inhibitors or ATRi could spill over and block ATM. To assess ATM activity, we monitored the appearance of phosphorylated serine 1981 (P-S1981) on ATM itself, a modification produced by ATM autophosphorylation after activation by DNA ends [25, 26]. For this we added dsDNA (8.9 kb) that contained a biotin group on one end so that we could recover the DNA back out of the extract and monitor the phosphorylation status of S1981 on DNA-bound ATM. The DNAs were added to HSS, and after incubation, they were isolated using magnetic streptavidin beads, washed, and bound proteins eluted. As shown in Figs. 1E&F, DNA-bound ATM contained the P-S1981 modification when the samples were treated with vehicle (DMSO), but not when ATMi was included. Notably, the appearance of P-S1981 was not impeded by ATRi, nor was it impacted by either DNA-PKcs inhibitor. Taken together, the data in Fig. 1 show that all the chemical inhibitors to be employed in this study are specific for their targets and there is no spill-over activity against the other kinases.

### DNA-PKcs is required for DSB-induced ATR signaling and represents a pathway running in parallel to MRN

Our previous work had identified at least two pathways that activate ATR in response to DSBs, one dependent on MRN and ATM and the other not [16]. It was conceivable that the ATM/MRN-independent pathway might represent events occurring during NHEJ-mediated repair of DSBs. Having established the specificity of the DNA-PKcs inhibitors, we therefore asked if blocking DNA-PKcs would impact ATR signaling. NU7441 was tested first, and we found that in a time-course experiment it efficiently blocked end-joining, as expected (Fig. 2A). Using the same samples, we also examined ATR signaling, with the appearance of P-CHK1 as the readout and found that the presence of NU7441 dampened but did not eliminate ATR signaling (Fig. 2D, compare lanes 1-3 with lanes 7-9). Similar results were obtained with AZD7648 (Fig. 2C and 2D, compare lanes 1-3 with lanes 10-12 in 2D). As expected, inclusion of ATRi efficiently blocked ATR signaling (Fig. 2D, lanes 4-6). These results thus link DNA-PKcs activity to ATR activation at DSBs.

**Figure 2:**
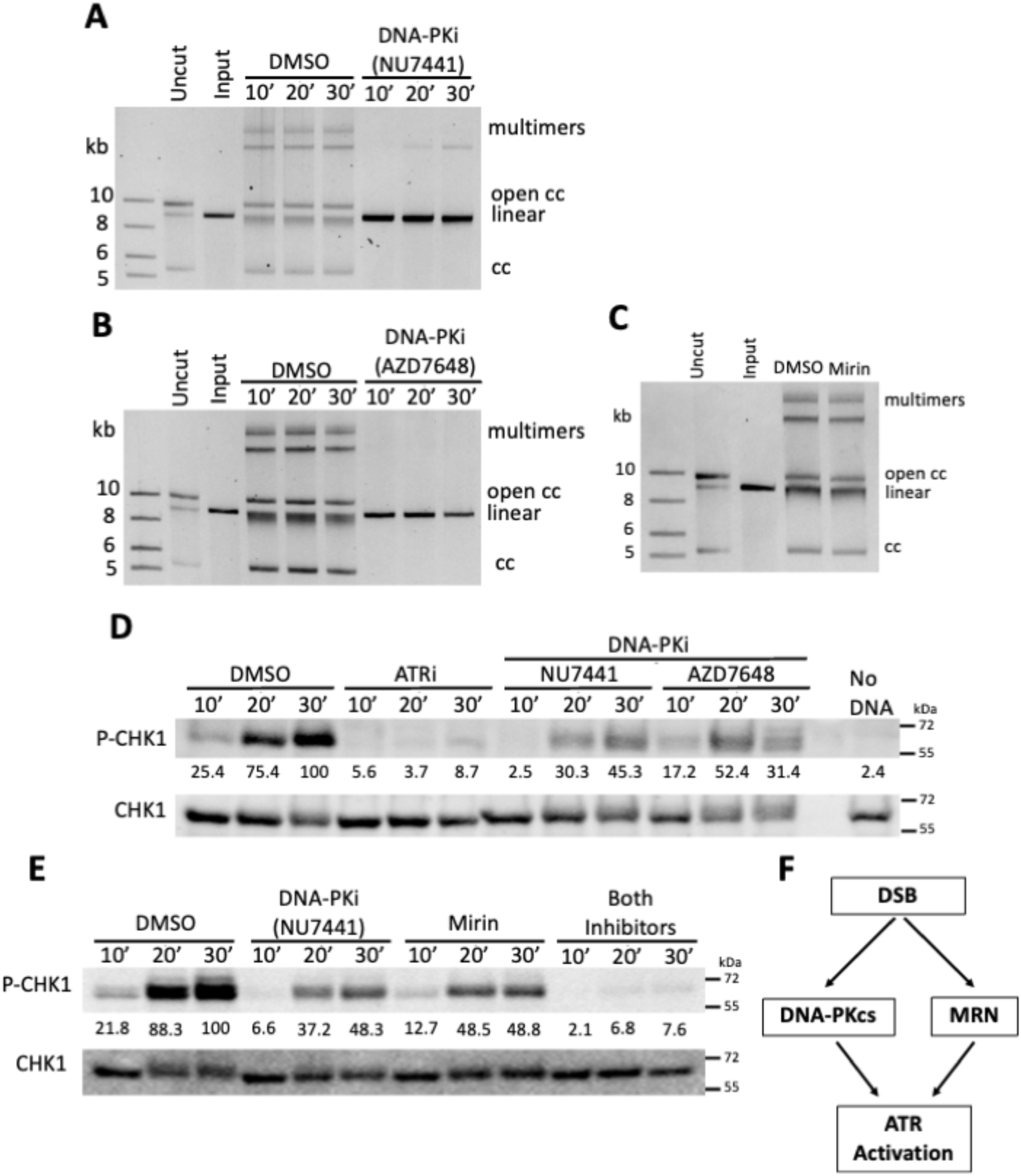
DNA-PKcs inhibition blocks NHEJ repair and suppresses ATR activation. A, a representative end-joining assay assessing NHEJ repair efficiency upon DNA-PKcs inhibition. XEE was treated with DMSO or a DNA-PKcs inhibitor (DNA-PKi; 100 µM NU7441). The XEE mixture was combined with the energy mix and the DSB substrate. After 10, 20, or 30 minutes of incubation, samples were digested with RNase A and Proteinase K and subjected to 0.6% agarose gel electrophoresis, followed by post-staining with SYBR Green. B, a representative end-joining assay examining NHEJ repair efficiency. The experiment was performed exactly as described in panel A. C, a representative end-joining assay testing NHEJ repair efficiency when MRE11, a subunit of the MRN complex, was inhibited by Mirin (100 µM). The experiment was carried out as described in B, except that the incubation was 30 minutes. D, a representative DMAX assay examining the effects of DNA-PKcs inhibition on ATR activation by DSBs. XEE was treated with DMSO, ATRi (100 µM Compound 45), or DNA-PKi (100 µM NU7441 [N] or 10 µM AZD7648 [A]). Samples received the DSB substrate (AfeI-digested 8.9 kb plasmid) and were incubated for 10, 20, or 30 minutes. After incubation, samples were subjected to Western blot analysis. E, a representative DMAX assay testing ATR activation when MRE11, DNA-PKcs, or both were inhibited. The experiment was performed as described in D. For D and E, total CHK1 served as a loading control. P-CHK1 signal intensity was quantified using ImageJ; the P-CHK1 signal from the DMSO-treated sample at 30 minutes was set to 100, and all other samples were normalized accordingly. Two biological replicates were performed, and a representative experiment is shown. Uncut, circular 8.9 kb plasmid; Input, AfeI-digested 8.9 kb plasmid, linear; DNA repair products: multimers, open circles, and closed circles (cc; supercoils). F, a schematic summarizing two parallel pathways - mediated by DNA-PKcs and MRN - for ATR activation in response to DSBs.

Our previous work had shown that loss of MRN would also dampen, but not eliminate, ATR signaling in the DMAX system. To pursue this we determined how mirin, a small molecule inhibitor of MRE11, impacted events in the DMAX system. There was no effect on NHEJ repair, as end joining occurred efficiently in the presence of 100 µM mirin (Fig. 2C). When ATR was examined, we found that mirin caused a partial reduction in signaling (Fig. 2E, compare lanes 1-3 to lanes 7-9). This partial reduction was similar to what we had previously reported with MRN depletion [16], and also echoed what was observed with the DNA-PKcs inhibitors (Fig. 2D&E). One explanation for these results is that MRN and DNA-PKcs control ATR activation pathways that run in parallel. If so, then we expected that ATR signaling would be suppressed to a greater extent when both mirin and NU7441 were combined, relative to when they are employed individually. As shown in Fig 2E, this is exactly what was observed as samples containing both inhibitors reduced ATR signaling to nearly undetectable levels. These data, taken together with our previous findings [16], show that two independently operating pathways, one led by DNA-PKcs and the other by MRN, act in parallel to activate ATR in the presence of DNA DSBs (Fig. 2F).

### NHEJ is dispensable for ATR signaling

Data presented thus far identify a role for DNA-PKcs in ATR activation by DSBs. This role could be direct, or indirect through DNA-PKcs’ role in promoting the NHEJ repair pathway. In other words, it may be that the process of NHEJ leads to ATR signaling in our system, and to pursue this, we chose to disrupt NHEJ independent of DNA-PKcs, via depletion of the core factors Ku or XLF. We produced antibodies targeting Ku80 and used them to remove the Ku complex from HSS via immunodepletion. As shown in Fig. 3A, Ku80 was readily detectable by Western blotting in a mock-depleted extract but not in the Ku80-depleted sample. We also prepared a recombinant version of Ku80 and added it back to the depleted HSS at approximately endogenous levels (Fig. 3A). Ku80 and Ku70 form an obligate heterodimer in egg extracts, and previous work has shown that depletion of one quantitatively co-depletes the other (17). We, therefore, also prepared a recombinant version of Ku70 and added it to the reconstituted extract. We then used these extracts to perform two assays, one an end-joining assay and the other an ATR activation assay. For end-joining, we found that loss of Ku caused an elimination of the closed circle repair product and severe diminishment of the multimer repair products, and this was rescued by addition of recombinant Ku (Fig. 3B). Importantly, unlike end-joining, ATR signaling occurred unimpeded in the Ku-depleted sample, while still being sensitive to the presence of a DNA-PKcs inhibitor (Fig. 3C). Thus, ATR signaling does not require Ku.

**Figure 3:**
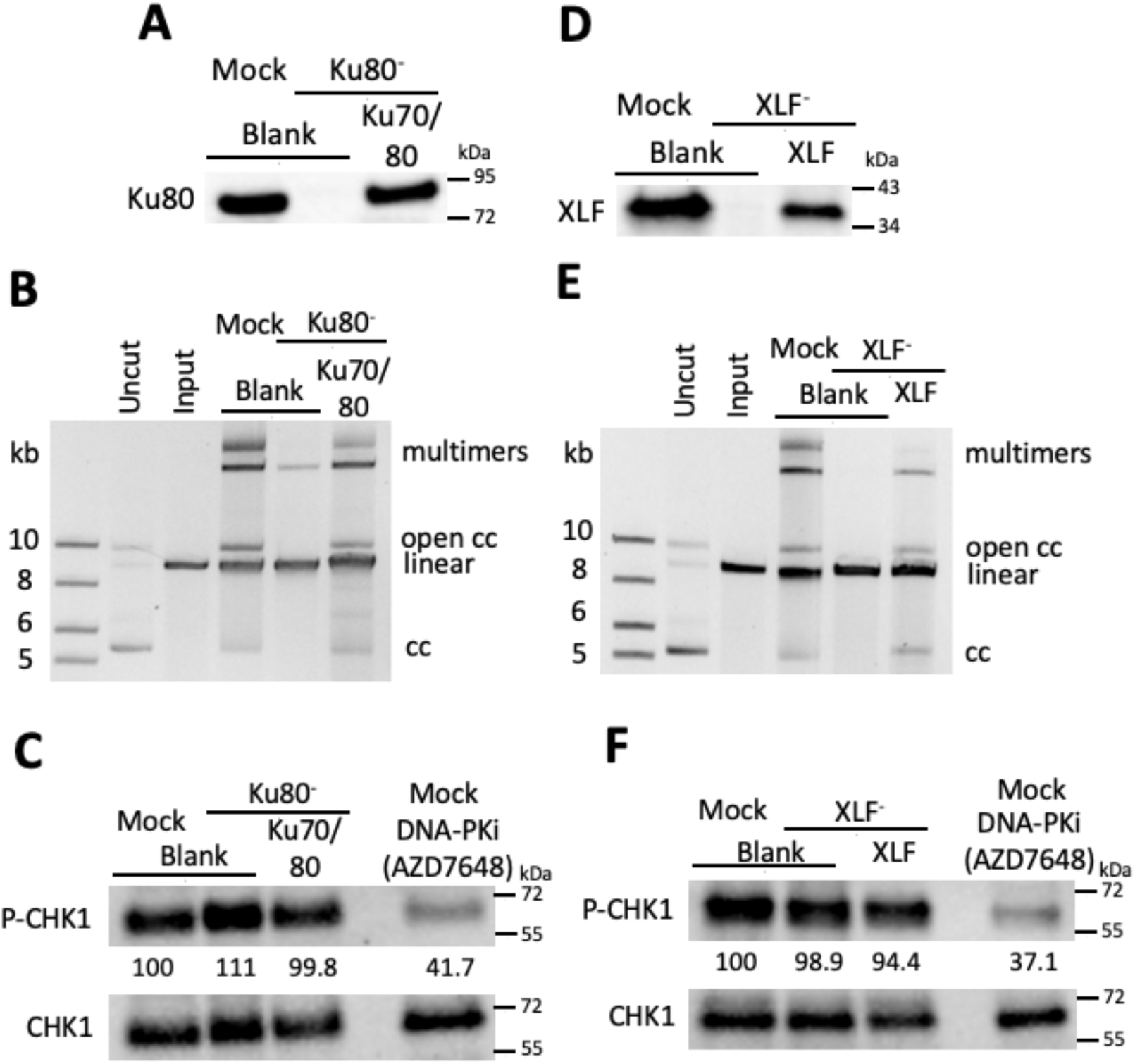
The Ku complex and XLF have no effect on ATR activation. A, a representative end-joining assay assessing NHEJ repair efficiency upon Ku complex depletion from XEE. XEE was mock-depleted or depleted of the endogenous Ku complex (consisting of the Ku70 and Ku80 subunits) and supplemented with unprogrammed IVTT (blank) or programmed IVTT to produce full-length Ku70 and Ku80. The XEE mixture was combined with the energy mix and the DSB substrate. After 30 minutes of incubation, samples were digested with RNase A and Proteinase K and subjected to 0.6% agarose gel electrophoresis, followed by post-staining with SYBR Green. B, a representative DMAX assay evaluating the effect of Ku complex depletion on ATR activation. XEE samples were prepared as described in A. An additional mock-depleted XEE sample was treated with DNA-PKi (AZD7648, 10 µM). All samples received the DSB substrate (AfeI-digested 8.9 kb plasmid). After 30 minutes of incubation, samples were subjected to Western blot analysis. C and D, a representative end-joining assay and a representative DMAX assay testing NHEJ repair efficiency and ATR activation, respectively, upon XLF depletion from XEE. The assays were performed exactly as described in A and B, respectively. Total CHK1 served as a loading control. P-CHK1 signal intensity was quantified using ImageJ; the P-CHK1 signal from the DMSO-treated mock sample was set to 100, and all other samples were normalized accordingly. Two biological replicates were performed, and a representative experiment is shown. Uncut, circular 8.9 kb plasmid; Input, AfeI-digested 8.9 kb plasmid, linear; DNA repair products: multimers, open circles, and closed circles (cc; supercoils).

We also raised antibodies against the core NHEJ factor XLF and used them to prepare XLF-depleted and reconstituted extracts (Fig. 3D). When these extracts were tested for end-joining we observed a severe defect that was rescued via addition of recombinant XLF (Fig. 3E). However, loss of XLF had no effect on ATR activation (Fig. 3F). Thus, like Ku, XLF is required for NHEJ but totally dispensable for ATR activation by DSBs. These results suggest that ATR signaling at DSBs is not interconnected with NHEJ and, in turn, that the role of DNA-PKcs in ATR activation can be separated from its role in promoting NHEJ.

### Ku-independent association of DNA-PKcs with DSBs promotes ATR signaling

The canonical model for NHEJ posits that DNA-PKcs is dependent on Ku for recruitment to DSBs (6). However, data from the Lieber group have shown that DNA-PK can bind DNA ends directly, in a manner where it retains catalytic activity [27, 28]. Thus, one possibility is that DNA-PKcs is present and active on DSBs even in the absence of Ku, as this would explain why ATR signaling readily occurs in the absence of Ku in a manner still dependent on DNA-PKcs (Fig. 3C). To pursue this, we performed a DSB binding assay, similar to what is shown in Fig. 1E, where dsDNA with a biotin group on one end is incubated in mock- or Ku-depleted extract and recovered on streptavidin beads. The beads were washed and eluted, and bound proteins were detected by Western blot. In this experiment we examined Ku, the ATR activator TOPBP1, DNA-PKcs, and XLF. As shown in Fig. 4A, all proteins were present in the total extract except Ku80, showing that neither DNA-PKcs, TOPBP1, nor XLF were co-depleted with Ku. When DSB-bound proteins were examined, we observed that both Ku and XLF were present in the mock-depleted sample, undetectable in the Ku-depleted sample, and present once again in the Ku80-depleted sample supplemented with recombinant Ku (Fig. 4A). This is consistent with the known requirement for Ku in recruitment of XLF to DSBs [29]. TOPBP1 levels on the DSB were unaffected by loss of Ku; however, we did observe a large decrease in DNA-PKcs levels on DSBs (Fig. 4A). Importantly, while the majority of DSB-bound DNA-PKcs requires Ku, a small amount can still associate with DSBs in the absence of Ku. Thus, it may be that this small, Ku-independent pool of DNA-PKcs is active in ATR signaling. To pursue this possibility, we performed an ATR activation assay with mock- or Ku-depleted extracts treated with the DNA-PKcs inhibitor AZD7648 and found that the inhibitor reduced ATR signaling to the same extent in both extracts (Fig. 4B). Thus, the small amount of DNA-PKcs on DSBs in the absence of Ku is sensitive to AZD7648, suggesting that it does indeed participate in ATR signaling. Based on these data, we propose that DNA-PKcs can work outside of NHEJ to promote ATR activation on DSBs (Fig. 4C).

**Figure 4:**
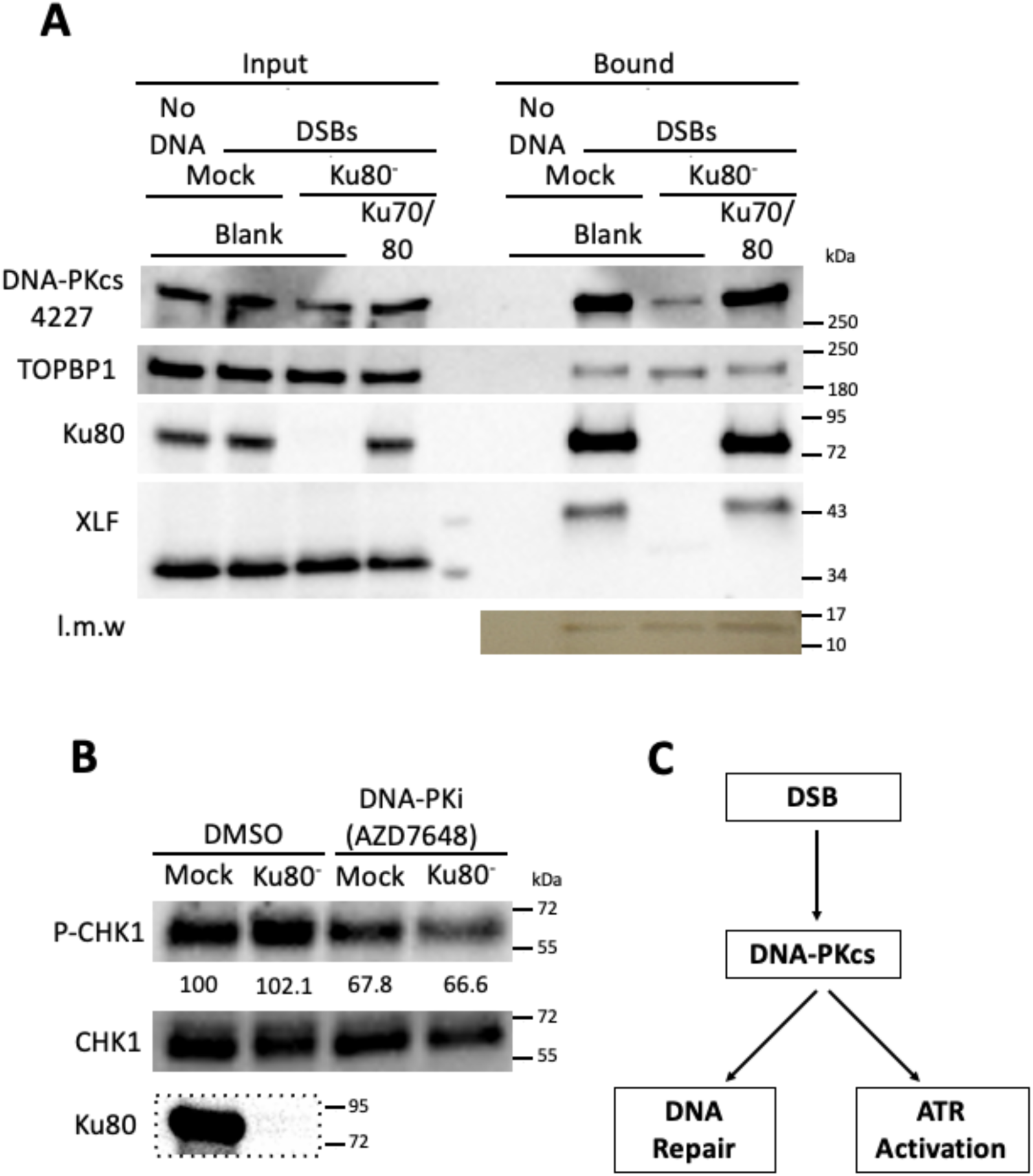
Ku-independent recruitment of DNA-PKcs to DSBs. A, a representative DSB-binding experiment determining the binding of DNA-PKcs, Ku80, XLF, and TOPBP1 to DSBs upon depletion of the Ku complex from XEE. XEE was mock-depleted or depleted of the endogenous Ku complex and supplemented with unprogrammed IVTT (blank) or programmed IVTT to produce full-length Ku70 and Ku80. Mock-depleted XEE was incubated with empty streptavidin beads (no DNA). The other three conditions (mock + blank, Ku80⁻ + blank, and Ku80⁻ + Ku70/80) used streptavidin beads loaded with 8.9 kb biotinylated DSBs. After incubation, samples were subjected to Western blot analysis. “Input” refers to a sample of the total XEE mixture taken before the addition of the DSB beads. “Bound” refers to proteins bound to the DSB beads. A human DNA-PKcs antibody (clone 4427) was used to detect *Xenopus* DNA-PKcs (see Materials and Methods). A low-molecular-weight (l.m.w.) band served as a loading control. B, a representative DMAX assay assessing ATR activation. XEE was mock-depleted or depleted of the endogenous Ku complex and subsequently treated with DMSO or DNA-PKi (AZD7648, 10 µM). All samples received the DSB substrate (AfeI-digested 8.9 kb plasmid) and were incubated for 30 minutes. After incubation, samples were subjected to Western blot analysis. P-CHK1 signal intensity was quantified using ImageJ; the P-CHK1 signal from the mock-depleted, DMSO-treated sample was set to 100, and all other samples were normalized accordingly. Two biological replicates were performed, and a representative experiment is shown. C, a schematic summarizing the functions of DNA-PKcs in response to DSBs, particularly in NHEJ repair and ATR activation.

### Physical and functional interactions between DNA-PKcs and TOPBP1

Our previous work has shown that, in frog egg extracts, TOPBP1 is the sole activator of ATR at DSBs [13]. Given our finding that DNA-PKcs also plays a role in ATR signaling we asked if these two factors interact with one another. Standard co-immunoprecipitations were performed, using HSS, and we observed that TOPBP1 and DNA-PKcs could indeed coprecipitate, in a reciprocal manner (Fig. 5A). The ability of DNA-PKcs and TOPBP1 to interact prompted us to examine if DNA-PKcs could impact phosphorylation of TOPBP1’s S1131, a modification essential to ATR signaling at DSBs but not at stalled replication forks [12]. Before addressing this, we first confirmed whether S1131 is required for ATR activation in the DMAX system. To address this, we performed an ATR activation assay in which TOPBP1 was depleted from XEE and rescued with either wild-type TOPBP1 or S1131A TOPBP1, a phospho-deficient mutant. As shown in Fig. S2A, exogenously added wild-type TOPBP1 restored ATR activation to endogenous levels (compare lane 3 with lane 1), while S1131A TOPBP1 failed to do so (lane 4). To further investigate whether S1131 is necessary for TOPBP1 recruitment to DSBs, we performed a DSB binding assay. Recombinant wild-type or S1131A TOPBP1 was added to XEE, and TOPBP1 binding to DSBs was assessed. As shown in Fig. S2B, both wild-type and S1131A TOPBP1 were able to bind DSBs. Based on these data we conclude that S1131 is essential for ATR signaling in the DMAX system, as expected, and that it acts after TOPBP1 recruitment to DSBs to promote ATR activation, also as expected.

**Figure 5:**
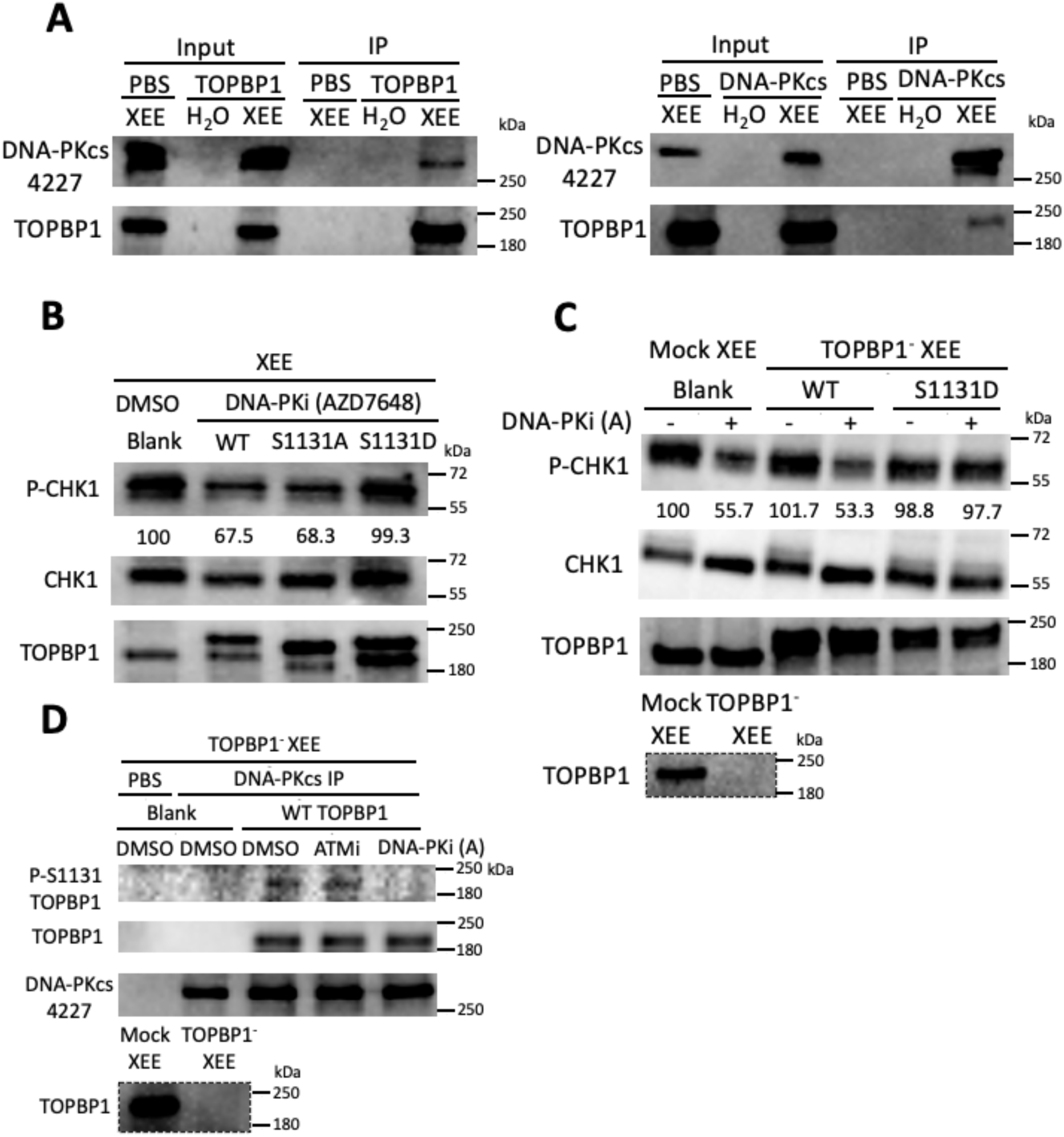
DNA-PKcs-mediated phosphorylation of TOPBP1 at Ser1131 promotes ATR activation. A, a representative reciprocal co-immunoprecipitation experiment determining whether TOPBP1 and DNA-PKcs interact in XEE. PBS, *Xenopus* anti-TOPBP1, or anti-DNA-PKcs antibodies coupled to Protein A beads were incubated with water or XEE. Samples were isolated from the beads and subjected to Western blot analysis. Because the TOPBP1 signal intensity in the first exposure was weak, a panel with a longer exposure is included for better visualization. B, a representative DMAX assay testing the ability of S1131 to override DNA-PKcs inhibition for ATR activation. Full-length wild-type (WT), S1131A, or S1131D TOPBP1, alongside unprogrammed IVTT (blank) as a control, were exogenously expressed via IVTT. These samples were added to XEE, which was subsequently treated with DMSO or DNA-PKi (10 µM AZD7648). All samples received the DSB substrate (AfeI-digested 8.9 kb plasmid) and were incubated for 30 minutes. After incubation, samples were subjected to Western blot analysis. C, a representative DMAX assay testing ATR activation. XEE was mock-depleted or depleted of endogenous TOPBP1 and supplemented with unprogrammed IVTT (blank) or programmed IVTT to produce full-length TOPBP1. The XEE mixture was then treated with DMSO or DNA-PKi (10 µM AZD7648). All samples received the DSB substrate (AfeI-digested 8.9 kb plasmid) and were incubated for 30 minutes. After incubation, samples were subjected to Western blot analysis. Total CHK1 served as a loading control. P-CHK1 signal intensity was quantified using ImageJ; the P-CHK1 signal from the DMSO-treated sample was set to 100, and all other samples were normalized accordingly. Two biological replicates were performed, and a representative experiment is shown. **D, a representative kinase assay examining TOPBP1 as a substrate of DNA-PKcs.** PBS or *Xenopus* DNA-PKcs antibodies coupled with Protein A beads were used to immunoprecipitate DNA-PKcs from mock-depleted XEE or XEE depleted of endogenous TOPBP1. Samples were then incubated with unprogrammed IVTT (blank) or programmed IVTT to produce wild-type (WT) TOPBP1, treated with DMSO, ATMi (100 µM KU60019), or DNA-PKi (10 µM AZD7648), and given the DSB substrate (AfeI-digested 8.9 kb plasmid). After 30 minutes of incubation, samples were subjected to Western blot analysis. Two biological replicates were performed, and a representative experiment is shown.

We next asked if a S1131D mutation in TOPBP1 could override the negative effects of DNA-PKcs inhibitors on ATR signaling. This mutation was chosen as previous work from the Dunphy group has shown that S1131D behaves as a phosphomimetic [12], and thus if the role of DNA-PKcs in ATR signaling is to promote P-S1131 then we expect that samples containing the S1131D mutant would be refractory to DNA-PKcs inhibitor. As shown in Fig. 5B, this is indeed the case, as extracts containing exogenous wild type TOPBP1, or a S1131A mutant, were sensitive to AZD7648 but the sample containing S1131D was not. To pursue these observations, we asked how the system would behave when S1131D was the sole source of TOPBP1 in the extract. As such, TOPBP1 was depleted and recombinant wild type or S1131D forms of TOPBP1 were added back. As expected, S1131D TOPBP1 overrode the suppression of the DNA-PKcs inhibitor for ATR activation (Fig. 5C). To further assess whether TOPBP1 is a substrate of DNA-PKcs, we performed an immunoprecipitation-kinase assay. As shown in Fig. 5D, wild-type TOPBP1 was phosphorylated after being incubated with DNA-PKcs immunoprecipitated extracts containing DSBs (lane 3). To rule out the possibility of co-immunoprecipitation of ATM in the extracts, we included two samples where the extracts were treated with ATM or DNA-PKcs inhibitor. Consistent with the absence of co-precipitated ATM, P-S1131 signal was unaffected by ATM inhibition (lane 4), but abolished by DNA-PKcs inhibition (lane 5). Taken together, these data suggest that DNA-PKcs targets and phosphorylates TOPBP1 at S1131.

### Both ATM and DNA-PKcs promote phosphorylation of TOPBP1 on S1131

Data shown thus far suggest a model whereby DNA-PKcs promotes ATR signaling at DSBs via phosphorylation of TOPBP1 on S1131. Previous work has shown that ATM can perform this same function, and thus one possibility is that the two ATR activation pathways we have identified in this work and previous work [16] boil down to the two kinases, ATM and DNA-PKcs, each phosphorylating this critical residue. To pursue this, we first asked if ATMi works additively with AZD7648 to reduce ATR activation in the presence of DSBs. As shown in Fig. 6A, this is indeed the case as addition of each inhibitor alone reduced the amount of P-CHK1 by ∼50% and when combined P-CHK1 was reduced to nearly background levels. We next looked directly at the S1131 status of TOPBP1 under these same conditions, using an antibody that specifically recognizes TOPBP1 P-S1131 [12], and observed that, like P-CHK1, P-S1131 was reduced by the individual treatments and reduced further by the combinatorial treatment (Fig. 6B&C). These data thus provide an explanation for the requirement for DNA-PKcs in ATR signaling at DSBs: the kinase promotes phosphorylation of TOPBP1’s S1131, leading to ATR activation.

**Figure 6:**
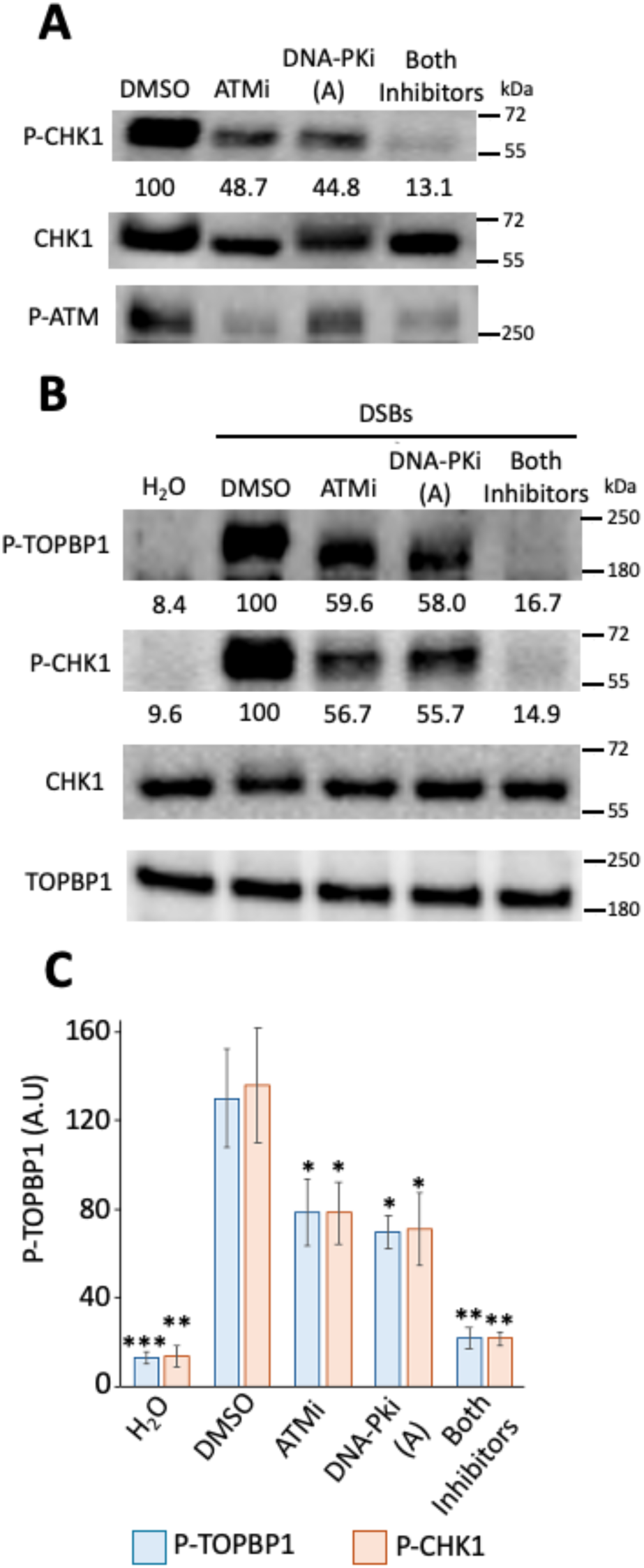
Phosphorylation of TOPBP1 at Ser1131 by ATM and DNA-PKcs. A, a representative DMAX assay testing the effects of inhibiting ATM, DNA-PKcs, or both kinases on ATR activation. XEE was treated with ATMi (100 µM KU60019), DNA-PKi (10 µM AZD7648 [A]), or both inhibitors and mixed with the DSB substrate (AfeI-digested 8.9 kb plasmid). After 30 minutes of incubation, samples were subjected to Western blot analysis. Two biological replicates were performed, and a representative experiment is shown. Total CHK1 served as a loading control. P-CHK1 signal intensity was quantified using ImageJ; the P-CHK1 signal from the DMSO-treated sample was set to 100, and all other samples were normalized accordingly. B, a representative DMAX assay testing the effects of inhibiting ATM, DNA-PKcs, or both kinases on phosphorylation of TOPBP1 at S1131 and on ATR activation. XEE was treated with ATMi (100 µM KU60019), DNA-PKi (10 µM AZD7648 [A]), or both inhibitors and mixed with water or the DSB substrate (AfeI-digested 8.9 kb plasmid). After 30 minutes of incubation, samples were subjected to Western blot analysis. Two biological replicates were performed, and a representative experiment is shown. Total TOPBP1 and CHK1 served as loading controls. P-TOPBP1 and P-CHK1 signal intensities were quantified using ImageJ; the P-TOPBP1 or P-CHK1 signal from the DMSO-treated sample was set to 100, and all other samples were normalized accordingly. C, a bar graph showing quantification of signal intensities of P-TOPBP1 and P-CHK1 in arbitrary units (A.U.). Error bars represent standard deviations; p-values were calculated using a t-test (n.s., not significant, p > 0.05; *, p < 0.05; **, p < 0.005; ***, p < 0.0005).

## DISCUSSION

The PIKK kinases - DNA-PKcs, ATM, and ATR - are the master regulators in responding to DNA damage [30]. DNA-PKcs and ATM are well-characterized for their roles in sensing DSBs and repairing them through either NHEJ or HR repair pathways. However, the mechanism for ATR activation at DSBs is poorly understood. Several studies have revealed context-dependent crosstalk among the three kinases and its significance in regulating DNA damage response pathways (Blackford and Jackson, 2017). For instance, ATM-dependent DNA-PKcs phosphorylation at Threonine 4102 facilitates NHEJ repair [31]. In addition, ATR directly phosphorylates DNA-PKcs upon replication stress [32]. Here, we report an uncharacterized role for DNA-PKcs in the DSB response, specifically in ATR activation. We find that this role is independent of DNA-PKcs’s canonical function in the NHEJ and is mediated by phosphorylation of the ATR activator TOPBP1 at S1131. Furthermore, this DNA-PKcs-dependent route operates in parallel with the previously described MRN/ATM-dependent pathway [16].

In this work, we identify DNA-PKcs as a key player driving the alternative pathway for ATR activation. The canonical pathway of ATR activation depends on MRN/ATM and is mediated by resection. While consistent with existing data, our findings establish an independent pathway that runs in parallel with the MRN/ATM-dependent pathway and is regulated by DNA-PKcs. This DNA-PKcs-dependent ATR activation is uncoupled from NHEJ repair, as loss of the Ku complex or XLF still leaves ATR activation intact (Fig. 3). Therefore, DNA-PKcs’s role is not a downstream consequence of the repair reaction itself. Since the Ku complex is dispensable for ATR activation, a direct implication of the NHEJ-independent pathway is that DNA-PKcs must be loaded on DSBs without Ku assistance. We find that a residual Ku-independent pool of DNA-PKcs remains associated with DNA ends in Ku-depleted extracts (Fig. 4A). This residual binding is consistent with biochemical work showing that purified DNA-PKcs can engage DNA ends directly in a manner that supports catalytic activity [27, 28, 33]. These findings help to mechanistically separate the functions of DNA-PKcs in NHEJ and ATR activation.

The mechanism by which DNA-PKcs promotes ATR activation appears to be phosphorylation of TOPBP1 on S1131, the same residue previously identified as an ATM substrate that is critical for ATR activation [12]. DNA-PKcs and ATM are known to share several substrates. A classic example is phosphorylation of H2AX at Ser139 by both kinases in response to ionizing radiation, with DNA-PKcs contributing particularly when ATM is compromised, thereby supporting chromatin remodeling and DNA repair [34]. Another is phosphorylation of p53 at Ser15 by both kinases following ionizing radiation, which activates p53’s role in programmed cell death [35, 36]. Here, we identify TOPBP1 as an additional substrate phosphorylated by both ATM and DNA-PKcs, and we show that phosphorylation at S1131 is crucial for activating ATR at DSBs. A previous study using human cell-free extracts reported that DNA-PKcs promotes phosphorylation of TOPBP1 at Ser1138 (the orthologue of *Xenopus* S1131) on a gapped DNA substrate mimicking a replication intermediate [37]. Our work extends this finding to DSBs, demonstrating that DNA-PKcs targets the same residue in response to a structurally distinct DNA lesion. Our work also extends this previous finding by showing that DNA-PKcs can work outside of NHEJ to promote P-S1131 on TOPBP1.

Neither DNA-PKcs nor ATM fully compensates for ATR activation when the other is inhibited, suggesting that pathways are isolated from one another (Fig. 6 and Fig.7). One plausible explanation lies in the differential accessibility of the MRN complex and DNA-PKcs to DSBs. The MRN complex recognizes DSBs through the C-terminal MRE11 DNA-binding domain and typically engages the adjacent DSB ends [38]. In the MRN-ATM-TOPBP1-ATR axis, NBS1 functions as a bridging factor that binds both ATM and TOPBP1 [15, 39], providing a defined route for substrate presentation. A notable feature of the DNA-PKcs-mediated pathway is that DNA-PKcs can bind DSB ends in the absence of the Ku complex. Early biochemical studies established that DNA-PKcs has Ku-independent DNA-binding activity [27, 28, 33], but this activity has received limited attention in the context of DSB signaling. Importantly, our data raise the possibility that the signaling and repair functions of DNA-PKcs are partitioned by the Ku complex: a Ku-loaded pool drives canonical NHEJ, whereas a Ku-independent pool contributes to ATR activation. It may also be the case that DNA-PKcs can engage TOPBP1 while bound to Ku, such that under normal conditions TOPBP1 is activated as a function of Ku recognition of DNA ends, and only in the absence of Ku does DNA-PKcs’ ability to bind DNA ends come into play. Overall, our finding that both DNA end recognition events, MRN/ATM for HR and Ku/DNA-PKcs for NHEJ, lead to activation of TOPBP1 at DSBs highlights the robustness of ATR activation as a safeguard for genome stability during DSB repair.

**Figure 7:**
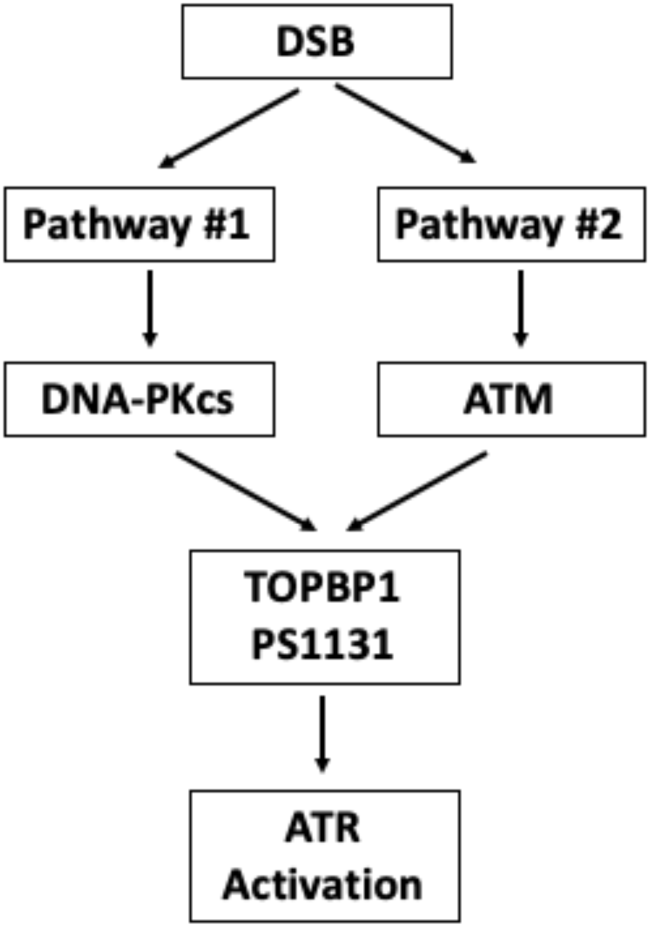
A schematic showing two distinct pathways (ATM-dependent and DNA-PK-dependent) for ATR activation at DSBs (refer to the Discussion for further details).

Our work settles the question of why TOPBP1 S1131 is absolutely required for DSB-based ATR signaling whereas MRN/ATM is only partially required as we have shown that DNA-PKcs is also a major player. Our work does however raise new questions. For example, in the DNA-PKcs dependent pathway, how is TOPBP1 recruited to DSBs? Our data show that DNA-PKcs and TOPBP1 can associate with one another in extracts lacking DNA, so it may simply be the case that DNA-PKcs carries TOPBP1 to the DNA end to initiate signaling. This is analogous to what occurs in the MRN/ATM pathway, where TOPBP1 binds to NBS1 and is thereby carried to DNA ends. A related and perhaps more interesting question is how, within the DNA-PKcs pathway, are ATR and ATRIP recruited to DSBs? We note that our experiments exclusively use a blunt-ended DNA substrate for ATR signaling, and this is unlikely to undergo extensive resection during NHEJ-based repair. As such, our data suggest an ssDNA-independent mechanism for ATR/ATRIP recruitment, and future experiments will address this exciting possibility. In summary, our work identifies DNA-PKcs as a mediator of ATR activation at DSBs, acting in parallel with the MRN/ATM pathway and converging on TOPBP1 S1131. These findings extend the functional repertoire of DNA-PKcs beyond NHEJ and refine current models of DSB-induced ATR activation.

## ACKNOWLEDGEMENTS

We thank Hovik Gasparyan, Shan Yan, Katrina Montales, and Kenna Ruis for construction of plasmids. We thank Hironori Funabiki for sharing the Ku70 and Ku80 expression vectors, Joseph Loparo for the DNA-PKcs expression vector, Michael Lieber for the DNA-PKcs 4227 antibody, and William Dunphy for the *Xenopus* P-S1131 TOPBP1 antibody. Grammarly was used to proofread and edit the manuscript.

## AUTHOR CONTRIBUTIONS

O.H.: Investigation, Data Analysis, and Visualization. O.H. and W. M. M: Methodology, Writing - original draft, review and editing. W. M. M: Conceptualization, Supervision, Project administration, Funding acquisition.

## SUPPLEMENTARY DATA

Supplementary Data are available at NAR online.

## DATA AVAILABILITY

All data are contained within the article.

