## Extended data for "An NHEJ-independent Role for DNA-PKcs in ATR activation at DNA Double-Strand Breaks"

### Supplemental Figures and Legends

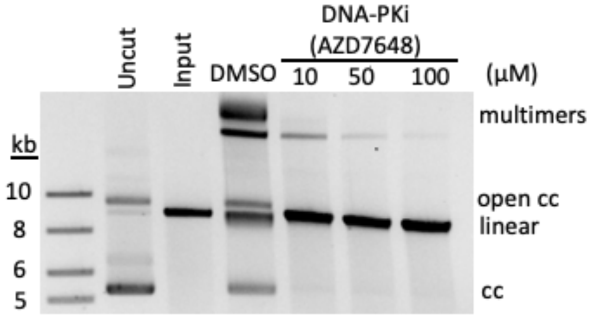

**Figure S1: Optimizing the working concentration of the DNA-PKcs inhibitor AZD7648**. A representative end-joining assay examining NHEJ repair efficiency when DNA-PKcs was inhibited by DNA-PKi (AZD7648) at different concentrations. XEE was treated with DNA-PKi at 10, 50, or 100 μM and mixed with the DSB substrate (AfeI-digested 8.9 kb plasmid). After 30 minutes of incubation, samples were digested with RNase A and Proteinase K and subjected to 0.6% agarose gel electrophoresis, followed by post-staining with SYBR Green. Uncut, circular 8.9 kb plasmid; Input, AfeI-digested 8.9 kb plasmid, linear; DNA repair products: multimers, open circles, and closed circles (cc; supercoils). Two biological replicates were performed, and a representative experiment is shown.

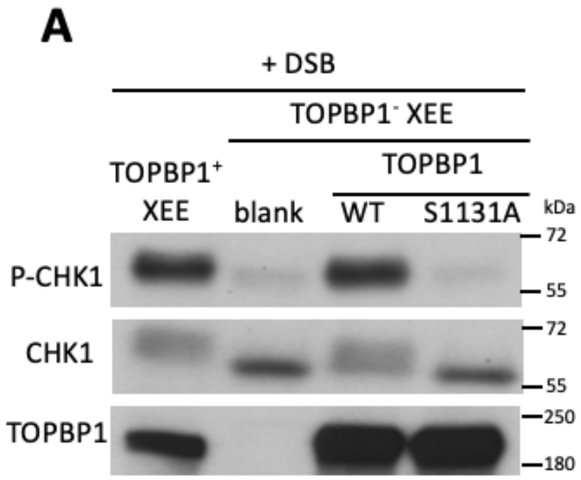
A
+ DSB
TOPBP1- XEE
TOPBP1
TOPBP1+ XEE
blank
WT
S1131A
kDa
P-CHK1
CHK1
TOPBP1
72
55
72
55
250
180
B
DSB binding assay
WT
S1131A
kDa
myc-TOPBP1 (bound)
myc-TOPBP1 (input)
l.m.w. protein (bound)
250
180
250
180
17
10

**Figure S2: S1131 in the AAD of TOPBP1 is dispensable for TOPBP1 recruitment to DSBs but is required for ATR activation in the DMAX system.** A, a representative DMAX assay examining ATR activation in XEE depleted of TOPBP1 and rescued with S1131A TOPBP1. XEE was mock-depleted or depleted of endogenous TOPBP1 and supplemented with unprogrammed IVTT (blank) or programmed IVTT to produce full-length wild-type (WT) or S1131A TOPBP1. The XEE mixture was combined with DSBs and incubated for 30 minutes. After incubation, samples were subjected to Western blot analysis. B, a representative DSB-binding assay testing the ability of wild-type (WT) or S1131A TOPBP1 proteins to bind to DSBs. Myc-tagged TOPBP1 proteins (full-length WT or S1131A mutant) were expressed via IVTT and added to XEE. The XEE mixture was combined with streptavidin beads loaded with biotinylated DSBs and incubated for 30 minutes. After incubation, samples were isolated from the beads and subjected to Western blot analysis. An anti-Myc antibody was used to probe for TOPBP1. A low-molecular-weight (l.m.w.) band served as a loading control.
